# Instantaneous isotropic volumetric imaging of fast biological processes

**DOI:** 10.1101/459370

**Authors:** Nils Wagner, Nils Norlin, Jakob Gierten, Gustavo de Medeiros, Bálint Balázs, Joachim Wittbrodt, Lars Hufnagel, Robert Prevedel

**Affiliations:** Cell Biology and Biophysics Unit, European Molecular Biology Laboratory, Heidelberg, Germany; Department of Medical Biochemistry and Biophysics, Karolinska Institute, Stockholm, Sweden; Centre for Organismal Studies, Heidelberg University, Heidelberg, Germany; Department of Pediatric Cardiology, University Hospital Heidelberg, Heidelberg, Germany; Developmental Biology Unit, European Molecular Biology Laboratory, Heidelberg, Germany; Epigenetics and Neurobiology Unit, European Molecular Biology Laboratory, Monterotondo, Italy

## Abstract

Capturing highly dynamic biological processes at sub-cellular resolution is a recurring challenge in biology. Here we show that combining selective volume illumination with simultaneous acquisition of orthogonal light-fields yields 3D images with high, isotropic spatial resolution and free of reconstruction artefacts, thereby overcoming current limitations of light-field microscopy implementations. We demonstrate Medaka heart and blood flow imaging with single-cell resolution and free of motion artefacts at volume rates up to 200Hz.

---

Many important tissue-scale biological processes occur in three dimensions and on millisecond time-scales, such as action potentials in neuronal networks, calcium signaling, or the dynamics of beating hearts in small animals. Investigating these phenomena requires imaging techniques capable of recording the intricate dynamics on a broad range of spatial and temporal scales. Several imaging techniques have been developed to address this challenge ranging from highly optimized point and line scanning^1,2^, to selective plane illumination^3^ or by reducing the dimensionality of the image acquisition^4–6^. While the former techniques illuminate, and capture, only a sub-volume at a time and thus sequentially record the volume, the highest speed can be achieved by illuminating and recording the entire volume at once.

Light-field microscopy (LFM) has emerged as a particularly elegant and powerful technique for fast and instantaneous 3D imaging in biology, as it captures a 3D image with a single camera snapshot. This is achieved by recording the entire 4D light-field, containing information about both the 2D location as well as the 2D angle of the incident light, on a single 2D camera sensor, from which the original 3D distribution of (fluorescent) emitters in a volume of interest can be deduced^7,8^. Since LFM offers volumetric imaging limited by the frame-rate of the camera only, it has predominantly found applications in areas where the combination of large field-of-view (FOV) and high speed are essential, such as the recording of neuronal activity across model organisms^9–12^. Despite its unique assets, current implementations of LFM suffer from a number of disadvantages which have compromised its full potential for effective 3D imaging: (1) Inherent trade-offs between spatial (lateral as well as axial) resolution and 3D FOV^7,8^; (2) a non-uniform spatial resolution across depth^8^ as well as (3) the strong presence of reconstruction artefacts around the nominal focal plane of the objective^8^, as well as at the edges of the 3D FOV due to fluorescence emitted from structures outside the volume of interest. Although these effects can in principle be mitigated to some extent by the incorporation of phase-masks in the optical imaging path^13^, the overall complexity combined with the generally low spatial resolution of previous LFM implementations, especially in the axial dimension, have so far prevented more wide-spread applications across biomedical research fields.

Here we present a new light-field microscopy scheme based on selective volume illumination and simultaneous recording of perpendicular light-fields (**Fig. 1a**) – termed Iso-LFM. By spatially confining the excitation light to the volume of interest we optimized signal-to-background contrast and minimized erroneous reconstruction artefacts from out-of-volume emitters which are otherwise excited in typical wide-field illumination schemes. Detecting the emitted fluorescence via two identical objectives placed at a perpendicular angle with respect to each other, and orthogonally to the illumination objective, further allowed us to perform dual-view data fusion and joint deconvolution of the simultaneously acquired light-fields. This effectively yielded an isotropic spatial resolution of ~2μm, a factor 2-8 better than previous LFM demonstrations. In addition to providing isotropic resolution, the dual-view capability of our microscope further minimizes resolution non-uniformity across the volume, an effect that is particularly prominent at image planes close to the border as well as the center of the volumetric FOV (V-FOV), and significantly reduces the presence of image planes containing reconstruction artefacts (so called artefact plane^8^). To demonstrate the system’s ability to capture fast 3D processes with isotropic resolution, we imaged the beating heart of a 8dpf Medaka larvae across a 300×300×300μm FOV at a volumetric speed of 143Hz. This allowed us to spatially and temporally resolve the myocardiac tissue layer with single-cell resolution, significantly better compared to traditional ‘single-view’ (epi-) light-field detection. Furthermore, we also demonstrate 3D imaging and tracking of blood flow in Medaka blood vessels with 200Hz volume rate, sufficient to capture the dynamics of individual blood cells without motion blur over 200×200×200μm.

**Figure 1.**
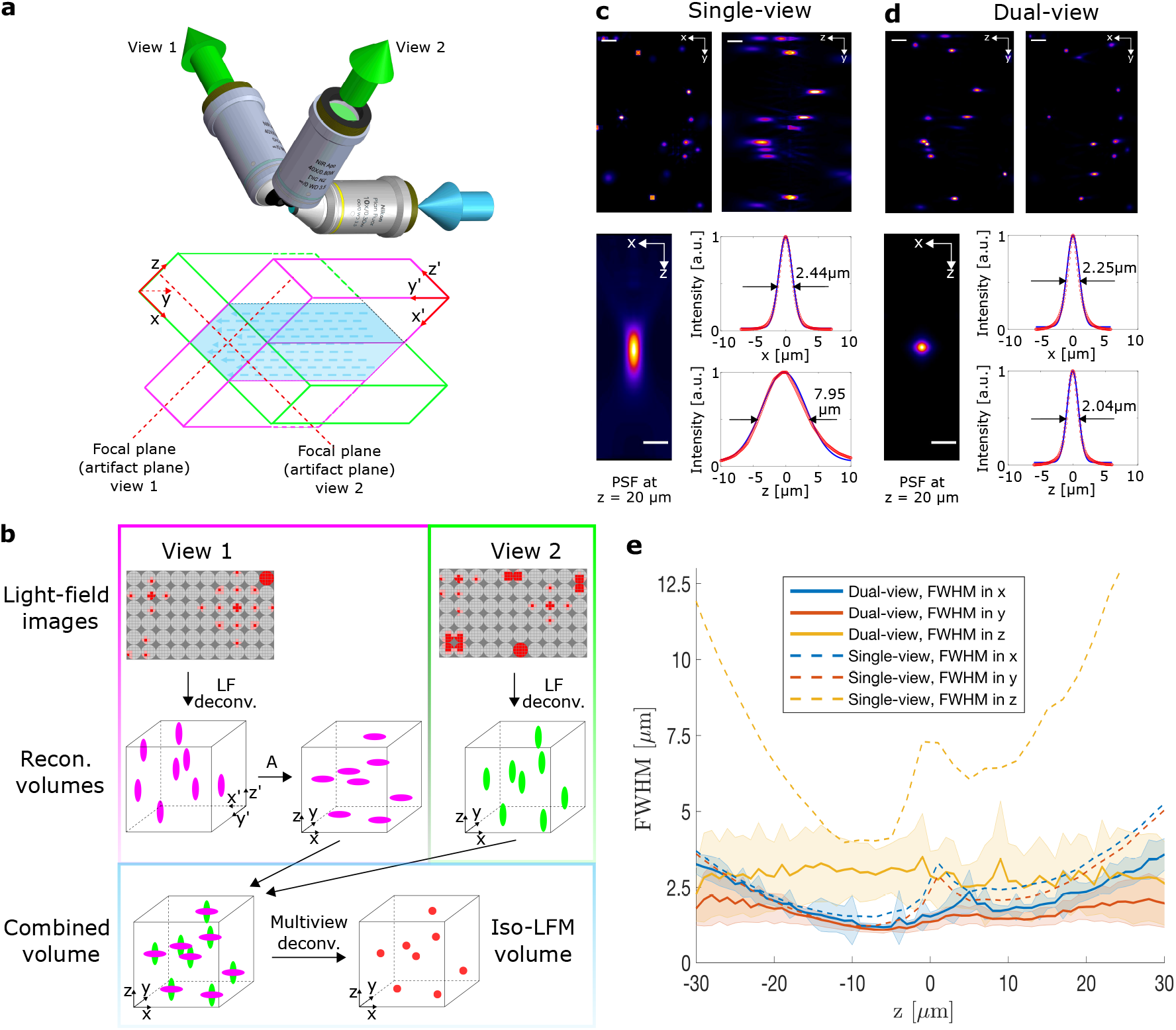
Principles and performance of Iso-LFM. (**a**) Objective configuration of Iso-LFM (blue, excitation; green, fluorescence detection). A schematic of the two individual light-field volumes is shown below, indicating the respective focal (artefact) planes as well as the overlapping volume (blue shading), (**b**) Iso-LFM reconstruction pipeline. Light-field images are individually reconstructed into 3D volumes, and an affine transformation A is applied to one volume to put both views in the same reference space. This allows subsequent multi-view deconvolution and results in isotropic resolution. Coordinate systems in (**b**) are in concordance with the coordinate systems in (**a**). (**c**) and (**d**): Maximum intensity projections of reconstructed fluorescent beads and their respective resolution (FWHM) for single-view and dual-view, respectively, demonstrating a ~3-fold improvement in axial resolution for Iso-LFM. The lower panels depict the measured PSF at 20 μm off the focal plane (averaged over n=10 beads) with lateral and axial FWHM indicated. Scalebars upper panel 10 μm and lower left panel 5 μm. (**e**) Mean values of lateral and axial PSF FWHM for the single-view and the dual-view case across imaging volume. Multiview deconvolution improves axial resolution across the entire overlapping volume and achieves near-isotropic resolution of ~2.0 ±0.8μm and ~2.9±1.2μm in x/y and z, respectively.

The mechanical design of the detection part of our Iso-LFM is based on an upright two-objective SPIM configuration^14^ with microlens arrays inserted in both detection pathways (**Online Methods** and **Supplementary Fig. 1,2**). To simultaneously introduce selective volume illumination for both objectives, a third objective is incorporated at an orthogonal angle. Employing two 40× 0.8NA detection objectives gives access to two imaging volumes of 300×300×60μm at perpendicular orientations with their intersecting volume (60×60×300μm) defining the volumetric FOV (V-FOV) of our Iso-LFM (**Fig. 1a**), while lower magnification objectives can significantly increase this V-FOV (e.g. 22.5× 0.5NA yield 300×300×300μm). To spatially combine the two simultaneously acquired light-field views into a single 3D volume with improved resolution, we implemented an automatic reconstruction pipeline that included image cropping, light-field resampling (‘rectification’), light-field deconvolution, affine volume registration in a common coordinate system as well as multi-view deconvolution^15^ with minimal to no user-intervention (**Fig. 1b, Online Methods, Supplementary Figure 3,4, Supplementary Note 1** and **Supplementary Software**). It is important to note, that due to the simultaneous illumination and acquisition of both light-fields no motion blur artefacts occur by combining both views. This is of particular importance when imaging very fast processes such as the *in-vivo* blood flow, where the time delay between sequentially acquired views would otherwise cause significant motion artefacts.

To evaluate the performance of our Iso-LFM, we imaged sub-diffraction sized beads suspended in agarose (**Fig. 1c,d**) and quantified the improvement in axial resolution owing to the dual-view (DV) capability compared to what could be obtained with a standard single-view (SV) configuration (**Fig. 1e**). Experimentally, we found that lateral and axial resolutions were almost identical (x,y ~2.0±0.8μm vs. z~2.9±1.2μm) and more uniform across imaging depth, a significant improvement compared to the SV-LFM case (average values x,y ~2.9±1.3μm and z~8.0±3.1μm, with x,y~5μm and z~15μm at the outer edges of the axial FOV). We also confirmed that the fluorescence of individual beads was assigned to the correct location throughout the volume, as verified by independent light-sheet imaging of the same volume. Furthermore, ‘tile-like’ artefacts close to the objectives focal plane that are common in standard LFM implementations were almost completely removed. Also, due to the selective volume illumination, we avoid background generated by fluorescent features outside the volume of interest.

In order to verify the suitability of Iso-LFM for capturing highly dynamic cellular processes in live animals, we performed *in-vivo* imaging of the beating heart in the juvenile Medaka fish (**Fig. 2a**). Employing two 20× (0.5NA) objective, we imaged cardiomyocytes labeled with a nuclear eGFP (*myl7⸬H2B-eGFP*) over a ~300×300×300μm V-FOV at a 143Hz volume rate, encompassing the entire heart (**Fig. 2b,c** and **Supplementary Video 1**). The spatiotemporal resolution of our Iso-LFM was sufficient to visualize the cardiomyocyte nuclei during the natural heart beat with single-cell resolution and free of reconstruction artefacts (**Fig. 2c**). In contrast, SV-LFM both showed cases of ambiguous cellular resolution as well as severely compromised image quality around the focal plane (**Fig. 2b**), a known limitation of SV-LFM. Furthermore, our Iso-LFM allowed us to track the rapid dynamics of blood flow in the cardiovascular system of Medaka with minimal motion blur (**Fig. 2d-f, Supplementary Figure 5,** and **Supplementary Video 2**). Here the instantaneous and high-speed nature of our microscope was essential to track undiluted eGFP-labelled circulating blood cells (*ENH-HMMA-4s-hsp70⸬eGFP*), due to the fast flow dynamics (up to 1400μm/sec) in these vessels. We note that conventional fast imaging approaches, including earlier LFM demonstrations^9^ based on a single-view would be unable to resolve these dynamics on the single-cell level.

**Figure 2.**
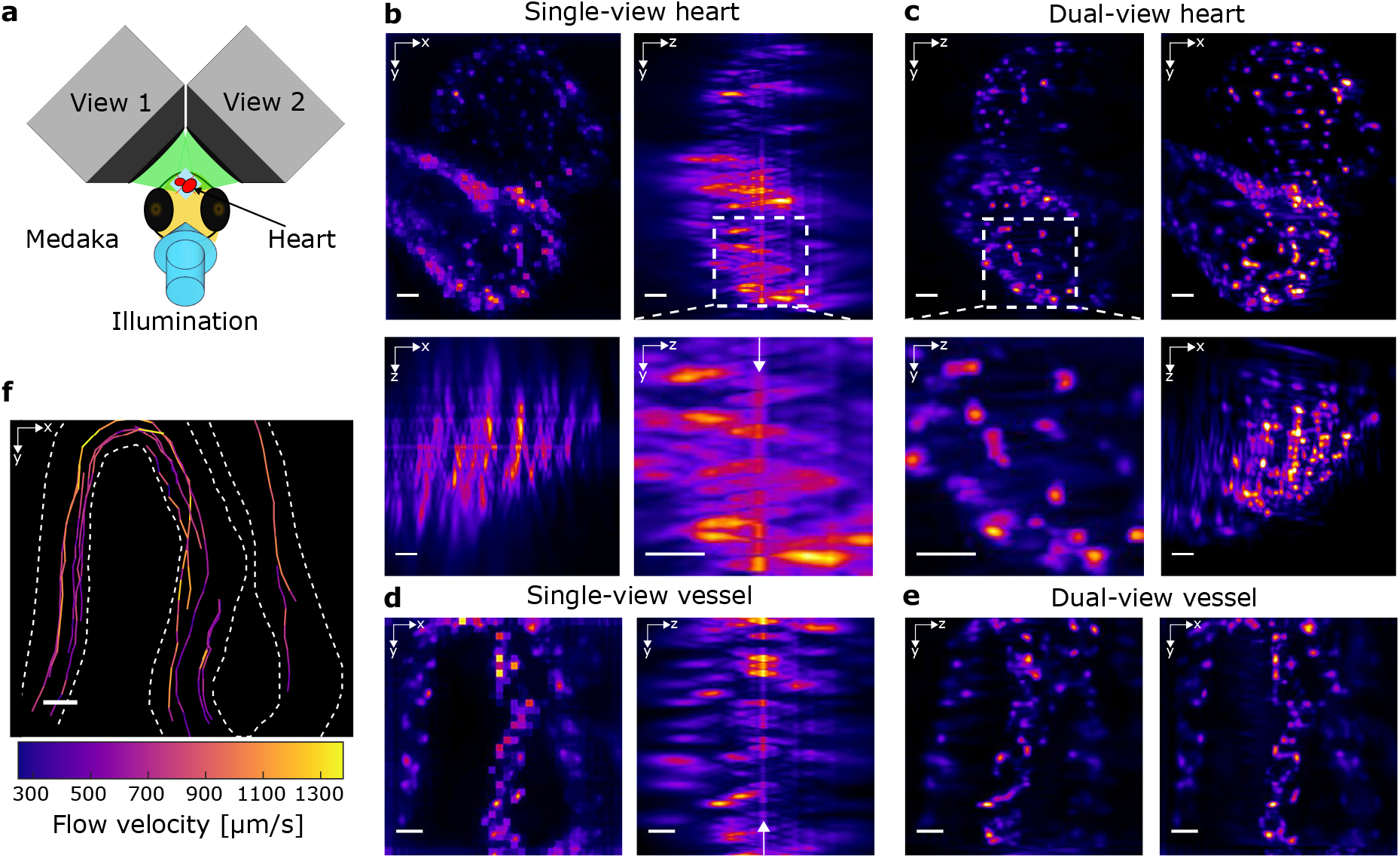
Instantaneous volumetric imaging of heart beating and blood flow in the juvenile Medaka fish using Iso-LFM. (**a**) Overview of the imaging configuration in Medaka. (**b**) Maximum intensity projections of single-view light-field images of the heart (*myl7⸬H2B-eGFP*), at 144Hz volume rate over 300×300×300μm. Low resolution and prominent artefacts are apparent (white arrow). (**c**) With dual-view Iso-LFM, single-cell resolution is clearly restored, and the artefact plane removed (c.f. zoom-ins denoted by white dashed boxes). Also see **Supplementary Videos 1,2**. (**d,e**) Blood flow imaging at 200Hz volume rate over 200×200×200μm. Dual-view Iso-LFM in (**e**) yields superior image quality compared to single-view light-field reconstruction (**d**) and enables tracking of individual blood cells at up to 1400μm/sec shown in (**f**). Also see **Supplementary Figure 5**. Scale bars 20 μm.

Previously developed microscopes, mostly based on light-sheet approaches, have also attempted to image fast biological processes such as the beating heart in the zebrafish larvae, e.g. by utilizing poor axial sampling combined with very short exposure times (and high excitation intensities) in order to rapidly acquire the required 3D volume^16^. Other demonstrations^17^ resorted to schemes in which the 3D heart was reconstructed from images acquired from different heart beats. This, however, can lead to volumetric ambiguity as the healthy heart is known to exhibit significant heart rate variability, thus making this approach less useful for meaningful cardiovascular investigations. Since gating of image acquisition to the cardiac cycle^18^ will also be sensitive to morphological fluctuations from heart beat to heart beat, a completely unbiased, direct volumetric imaging modality at minimal excitation intensity is still a crucial prerequisite for studying the dynamic properties and functions of the cardiovascular system with single-cell resolution. Therefore, Iso-LFM could become an important tool for studying dynamics on these fast timescales, potentially giving new mechanistic insights into medically relevant dysfunctions of the heart and blood transport system.

In summary, we have implemented a selective volume illumination dual-view light-field microscope and demonstrated its ability to image highly dynamic processes with sub-cellular isotropic resolution and minimal artefacts. The increased and more uniform spatial resolution together with higher contrast were crucial for imaging the beating heart and blood flow dynamics of 8dpf Medaka at single-cell resolution and with 200Hz volume rate. The simultaneous use of two objectives in Iso-LFM effectively doubles the light collection efficiency and eliminates spatio-temporal ambiguity associated with 3D imaging approaches based on sequential recording of image planes. Currently the upper limit in imaging speed is given by the camera read-out time (4ms) and is not fluorescence limited (exposure time 1ms). Thus, we foresee that future camera developments featuring multi-area read-out architectures can further increase the overall imaging speed towards the volumetric kHz regime. This would make Iso-LFM an attractive tool to record voltage dynamics^19^ on millisecond time scales in 3D within small neuronal networks. While the restricted V-FOV is a current drawback of Iso-LFM and thus requires proper sample positioning for the volume of interest, we also note that this approach scales favorably for lower objective magnifications and therefore V-FOVs (**Supplementary Note 2**). This is because the achievable (isotropic) resolution in multi-view deconvolution is predominantly determined by the lateral extent of the light-field PSF, which is less affected by lower objective magnifications^8^. We further note that our scheme is complementary to, and compatible with, standard strategies to obtain larger V-FOV, such as image tiling via sample movement or other means to remotely reposition the detection volumes^4,20^. Future work will focus on a unified and more efficient computational reconstruction scheme, including deep learning^21^, as well as the use of simplified objective geometries^22^ that would enable extensions to mammalian tissues^11^. Finally, we expect that future developments in camera sensor technologies and processor speed will contribute to further advances of Iso-LFM as well as other microscopy approaches based on the light-field concept.

## Acknowledgments

We would like to thank the EMBL Heidelberg mechanical and electronic workshop for help as well as the IT-Department for HPC cluster support. We further thank T. Thumberger for advice on CRISPR/Cas donor design and M. Majewsky, E. Leist and A. Saraceno for fish husbandry. J.G. is a fellow of the MD/PhD program of the Medical Faculty Heidelberg and of the Heidelberg Biosciences International Graduate School (HBIGS) and is grateful to M. Gorenflo for supervision and guidance. J.G. was supported by a Heidelberg Research Center for Molecular Medicine (HRCMM) Career Development Fellowship (CDF) and by the Deutsche Herzstiftung e.V. (S/02/17). N.N acknowledges support from VINNOVA and Åke Wiberg foundation. This work was supported by the European Molecular Biology Laboratory (N.W., N.N., B.B., G.M., L.H and R.P.).

## Author Contributions

N.N., L.H. and R.P. designed the imaging system. N.W. and N.N. built the imaging system and performed experiments together with J.G. and G.M. R.P. designed microlenses, J.G. generated transgenic animals under guidance of J.W. N.W. wrote analysis software with help from N.N. B.B. contributed to hardware control and data processing, and N.N. wrote computational cluster pipeline with assistance from N.W. R.P. and L.H. conceived and led the project.

## Online methods

### Iso-LFM optical setup

The microscope consists of one illumination and two opposing detection arms (see **Supplementary Fig. 1**). The illumination source is continuous-wave laser (λ = 488 nm, 50 mW, Omicron). We use a 10× 0.3NA (Nikon) water dipping objective for illumination and either a 40× 0.8NA (Nikon) or 20× 0.5NA (Olympus) water dipping objectives for detection. For the latter, our tubelens with focal length of 200mm yielded an effective magnification of 22.5×. Each detection arm features a microlens array (pitch 125 μm and focal length 3.125mm, RPC photonics) mounted in a 6-axis kinematic mount (Thorlabs, K6XS) allowing fine adjustment of the array in respect to the optics axis. The microlens array is imaged onto a 4.2 Megapixel (2048×2048 pixels) sCMOS camera (Andor Zyla) with a 1:1 relay macro lens objective (Nikon AF-S 105mm 2.8 G VR IF-ED Micro). The selective illumination is generated using a variable beam size illumination through a lens focusing on a relayed back aperture plane. For rapid imaging (>50 Hz) of juvenile Medaka we used a rod-like illumination. For this the laser beam was expanded and only the central plateau of the Gaussian illumination profile was used to illuminate the entire V-FOV simultaneously. The lateral extensions of the V-FOV could be adapted by changing the size of a square-shaped aperture that was used to crop out the central region of the laser beam. With this configuration, we were able to image with a 1 ms exposure time. Camera readout times as low as 4 ms could be achieved by cropping the sensor readout size. The sample stage consisted of a composite XYZ linear positioning stage (Newport M-562-XYZ) together with a piezo stage (Nanos LPS-30-30-1-V2_61-S-N and controller MC101) and a small rotation stage (Standa, 7R128), which together enabled precise positioning of the Medaka fish for imaging. Further information on the setup and can be found in **Supplementary Figs. 1,2**.

### Light-field deconvolution

Light-field reconstructions were conceptually based on previously published light-field deconvolution schemes^8^,^9^, which we further automatized and customized for our Iso-LFM implementation. First, the light-field point spread functions were calculated from vectorial diffraction theory analogous to Ref. ^9^. Then, automated custom written scripts were used to minimize human involvement and potential error in the subsequent reconstruction steps (see **Supplementary Figs. 3,4** and **Supplementary Software**). Timepoints were saved as single images that were then further distributed for deconvolution on a computation cluster platform (EMBL Heidelberg computational cluster https://wiki.embl.de/cluster/Hardware). The details of the reconstruction pipeline are further discussed in **Supplementary Note 1**. Reconstruction times depended on several parameters such as image size, number of deconvolution iterations, reconstruction volume, type of reconstruction method, and computational resources. Typically, utilizing our cluster pipeline work flow we managed to perform up to ~3000 volume reconstructions within 24h (~125/hour), a factor of ~20-40 times faster than GPU deconvolution on a dual-CPU workstation with 128GB memory and graphics card (Nvidia GTX 1070).

### Orthogonal light-field fusion

We implemented a semi-automated fusion pipeline to spatially combine the simultaneously acquired light-field images into a single 3D volume with improved spatial resolution. In order to find the correct affine transformations for registering the orthogonal views, we imaged a three-dimensional distribution of 0.1-μm fluorescent beads (TetraSpeck, Thermo Fisher Scientific) in agarose. The individually reconstructed volumes of the two views were then segmented and registered using the “Multiview-Reconstruction” Fiji-Plugin^15^. The affine transformations found during this procedure were subsequently used as a template and applied to each timepoint of the Iso-LFM recording. The volumes were then fused to obtain isotropic spatial resolution using the same Fiji-Plugin with the PSFs for multiview-deconvolution extracted from the bead image data. More details on the reconstruction pipeline are provided in **Supplementary Note 1**.

### Spatial resolution analysis

To quantify the performance of our setup we imaged a three-dimensional distribution of 0.1-μm fluorescent beads (TetraSpeck, Thermo Fisher Scientific) in agarose and the FWHM of the reconstructed beads was used as a resolution measure. Bead locations were detected with the “Multiview Reconstruction” plugin in Fiji. We then cropped a small volume around each detected bead position from the segmentation step in Fiji and fitted a Gaussian function to determine the FWHM using a custom written MATLAB script. Gaussian fits were performed as 1D line fits and 3D volume fits respectively. Both fitting schemes gave similar results. To compare single-view and dual-view data the same beads were used for the resolution measurement. Light-field deconvolution for these images was based on 8 iterations and yielded 31 distinct z planes, spaced 2 μm apart.

### Fish husbandry and transgenic lines

Medaka (*Oryzias latipes*) husbandry (permit number 35-9185.64/BH Wittbrodt) and experiments (permit number 35-9185.81/G-145/15 Wittbrodt) were performed according to local animal welfare standards (Tierschutzgesetz §11, Abs. 1, Nr. 1) and in accordance with European Union animal welfare guidelines. For *in-vivo* imaging, embryos were kept in 165 mg/l 1-phenyl-2-thiourea (PTU) in embryo rearing medium (ERM) from 1 dpf until imaging to inhibit pigmentation. The following transgenic lines were used: *ENH-HMMA-4s-hsp70⸬eGFP* labeling mature blood cells^23^ and *myl7⸬H2B-eGFP*. The *myl7⸬H2B-eGFP* transgene is contained in a PhiC31 landing site, which was modified based on a previous version^24^ and comprises a cassette of *FRT-myl7⸬HA-attP-H2B-eGFP-HA-pA-FRT3*. The modules of this cassette were cloned and assembled with the Golden Gate system^25^ and inserted into intron 1 of *nfia* (Ensembl ID ENSORLG00000003529.1) via homology-directed repair using CRISPR/Cas. Microinjections of donor plasmid, sgRNA and Cas9 mRNA into wild-type medaka Cab embryos at one cell-stage were performed as described earlier^26^.

### Medaka imaging

Medaka larvae were imaged 1-3 days after hatching. Hatchlings were anesthetized in 150 mg/l disodium phosphate-buffered (pH 7,3) tricaine and mounted in 1 % low-melting agarose (in ERM) containing 150 mg/l tricaine. For both imaging the beating heart as well as the blood flow, a light-field PSF was chosen that yielded 79 distinct *z* planes, spaced 4 μm apart after 8 iterations of deconvolution. Single blood cells were manually tracked in 3D using the Fiji plugin MtrackJ^27^. Calculated velocities and plots of the trajectories were done using Matlab 2018 (Mathworks).

### Code availability

All custom code used during the current study are either available as Supplementary Software or from the corresponding authors on reasonable request.

### Data availability

The datasets generated and/or analysed during the current study are available from the corresponding author on reasonable request.

